# Pathogenesis of Breakthrough Infections with SARS-CoV-2 Variants in Syrian Hamsters

**DOI:** 10.1101/2023.01.12.523876

**Authors:** Jessica Plunkard, Kathleen Mulka, Ruifeng Zhou, Patrick Tarwater, William Zhong, Margaret Lowman, Amanda Wong, Andrew Pekosz, Jason Villano

## Abstract

Severe acute respiratory syndrome coronavirus 2 (SARS-CoV-2), the causative agent of COVID-19, has evolved into multiple variants. Animal models are important to understand variant pathogenesis, particularly for those with mutations that have significant phenotypic or epidemiological effects. Here, cohorts of naïve or previously infected Syrian hamsters (*Mesocricetus auratus*) were infected with variants to investigate viral pathogenesis and disease protection. Naïve hamsters infected with SARS-CoV-2 variants had consistent clinical outcomes, tissue viral titers, and pathology, while hamsters that recovered from initial infection and were reinfected demonstrated less severe clinical disease and lung pathology than their naïve counterparts. Males had more frequent clinical signs than females in most variant groups, but few sex variations in tissue viral titers and lung pathology were observed. These findings support the use of Syrian hamsters as a SARS-CoV-2 model and highlight the importance of considering sex differences when using this species.

**Importance:** With the continued circulation and emergence of new severe acute respiratory syndrome coronavirus 2 (SARS-CoV-2) variants, understanding differences between the initial and a subsequent reinfection on disease pathogenesis is critical and highly relevant. This study characterizes Syrian hamsters as an animal model to study reinfection with SARS-CoV-2. Previous infection reduced the disease severity of reinfection with different SARS-CoV-2 variants.

## Introduction

SARS-CoV-2 variants are genetic lineages of the virus with mutations that alter virus replication, disease, transmission or evasion of adaptive immune responses (1). These were first noted in early 2020, only a few months after the first reported COVID-19 cases (2). Multiple genetically-distinct variant lineages have since emerged, and have been classified based on their genomic sequences via the Pango dynamic nomenclature system (3). An epidemiological classification system was also established to characterize variants with potentially deleterious human health consequences into three categories: Variants under monitoring (VUM), variants of interest (VOI), and variants of concern (VOC), with VOC having the most potential for impacting public health and disease outcomes (2). The WHO further labels individual VOC and VOI using the Greek alphabet (2).

Animal models have been used to study SARS-CoV-2 pathogenesis, transmission, vaccine candidates, and therapeutics (4). As with previous coronaviruses, Syrian hamsters stand out as an animal model (5). The virus replicates efficiently in their airways and nasal passages without progressing to severe disease in adult animals (6–8), modeling the outcome most seen in human patients. Here, the pathogenesis of SARS-CoV-2 variants in naïve or previously infected hamsters was investigated using clinical outcomes, lung histopathology, and respiratory tissue viral titers.

## Results

Table 1 identifies the isolate, Pango lineage, and WHO Greek alphabet designation (as applicable) for variants used in this study.

**TABLE 1.**
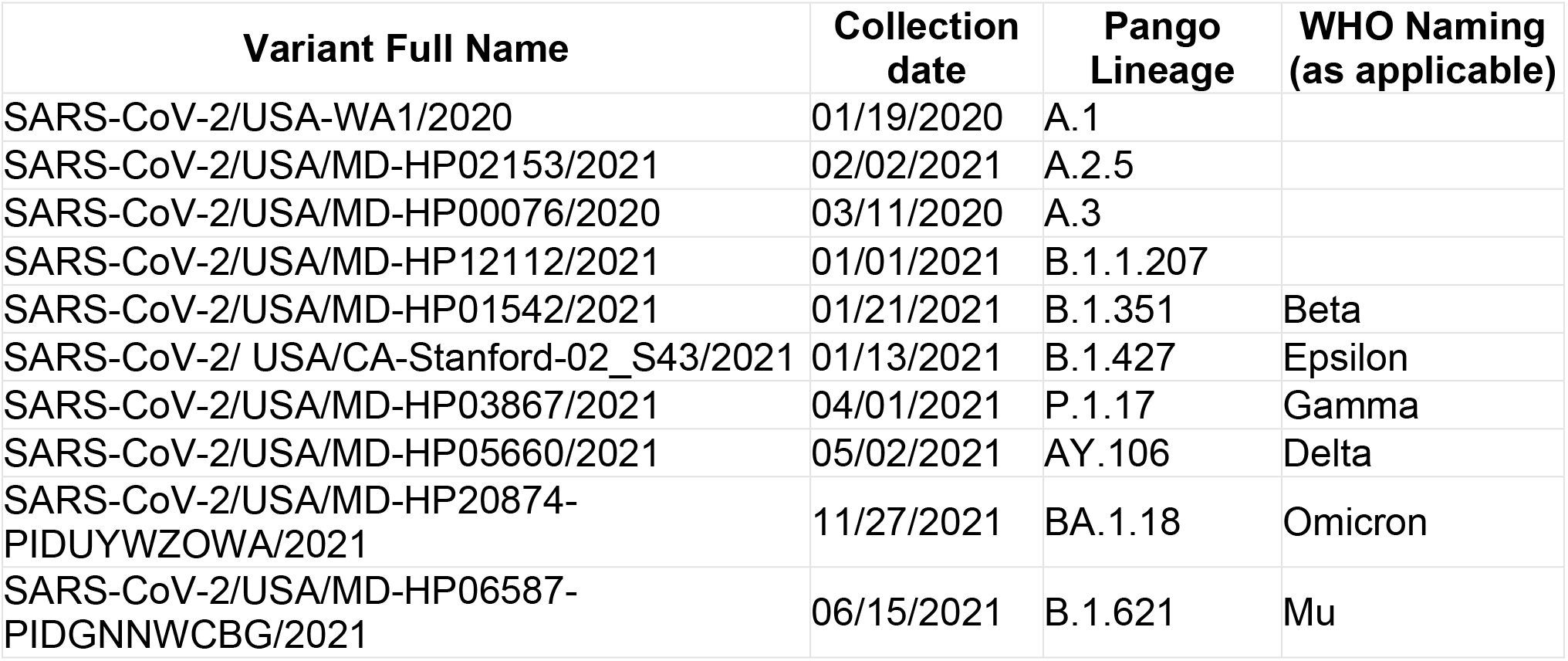
List of variants utilized.

| Variant Full Name | Collection date | Pango Lineage | WHO Naming (as applicable) |
| --- | --- | --- | --- |
| SARS-CoV-2/USA-WA1/2020 | 01/19/2020 | A.1 |  |
| SARS-CoV-2/USA/MD-HP02153/2021 | 02/02/2021 | A.2.5 |  |
| SARS-CoV-2/USA/MD-HP00076/2020 | 03/11/2020 | A.3 |  |
| SARS-CoV-2/USA/MD-HP12112/2021 | 01/01/2021 | B.1.1.207 |  |
| SARS-CoV-2/USA/MD-HP01542/2021 | 01/21/2021 | B.1.351 | Beta |
| SARS-CoV-2/ USA/CA-Stanford-02_S43/2021 | 01/13/2021 | B.1.427 | Epsilon |
| SARS-CoV-2/USA/MD-HP03867/2021 | 04/01/2021 | P.1.17 | Gamma |
| SARS-CoV-2/USA/MD-HP05660/2021 | 05/02/2021 | AY.106 | Delta |
| SARS-CoV-2/USA/MD-HP20874-PIDUYWZOWA/2021 | 11/27/2021 | BA.1.18 | Omicron |
| SARS-CoV-2/USA/MD-HP06587-PIDGNNWCBG/2021 | 06/15/2021 | B.1.621 | Mu |

### Naïve Infection with SARS-CoV-2 Variants

#### Clinical outcomes

Clinical outcomes were monitored daily to evaluate differences between variants and sexes. Hamsters lost body weight (BW) over the first 6 days post infection (dpi) and began regaining weight at 7 dpi, with some differences in mean BW appreciated. Groups infected with A.3 exhibited significantly more BW loss compared to A.2.5, B.1.1.207, Epsilon, Delta, and Omicron groups, losing up to 26% BW by 6 dpi. Omicron infected animals lost significantly less BW than A.3, Beta, and Gamma groups, with a maximal loss of only 6% BW (Fig. 1A). There was no significant sex-associated impact on BW.

**FIGURE 1.**
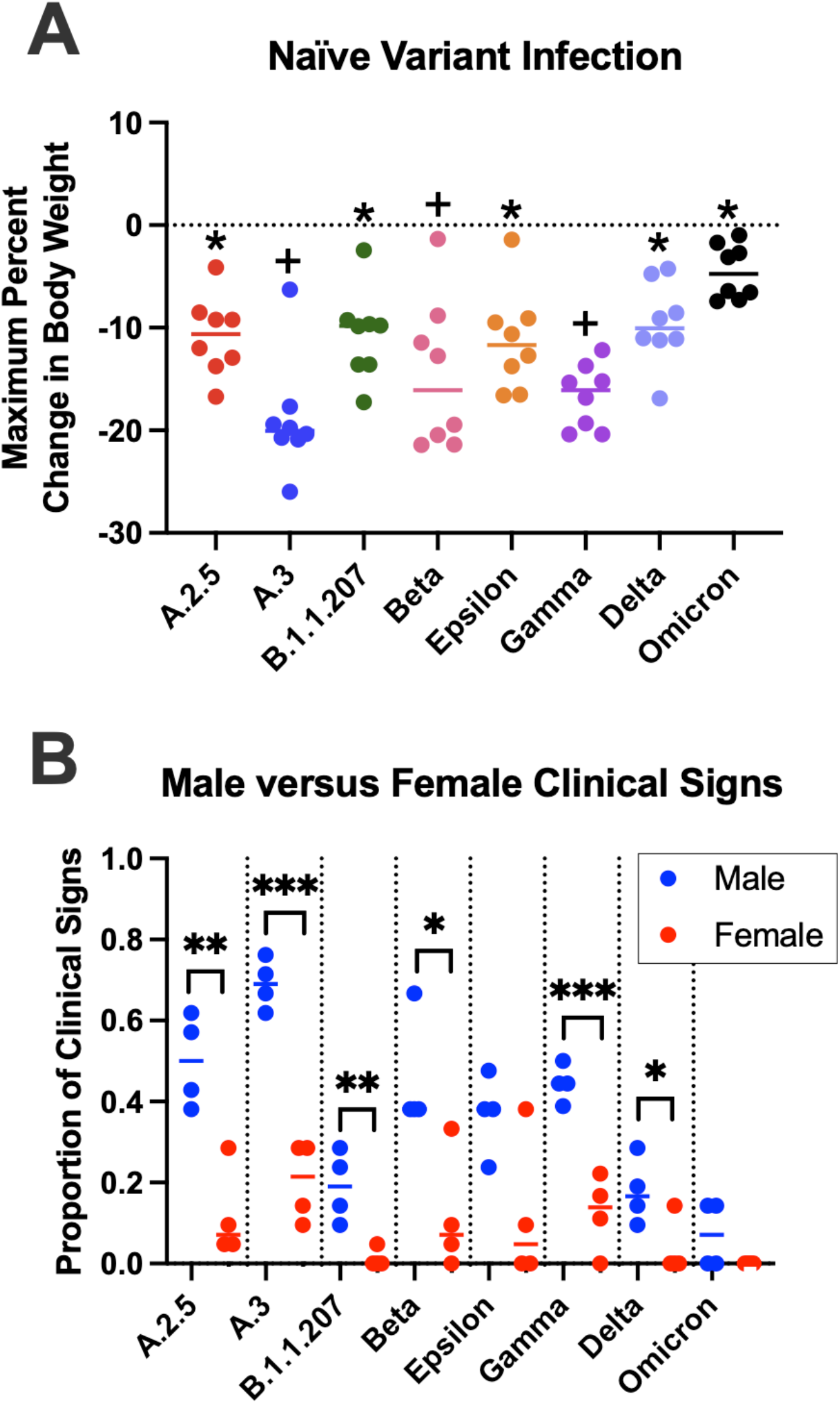
Clinical signs following naïve infection. (A) The maximum percent body weight change over 7 dpi. Using a one-way ANOVA, few significant differences in body weight changes were observed between variant groups, apart from the Omicron group experiencing significantly less weight loss than several groups (denoted with a +) and A.3 experiencing significantly more weight loss than several other groups (denoted with a *). (B) Although few sex differences were observed between variant groups for body weight change, significant differences were noted in the frequency of other clinical signs between all groups except Omicron and Epsilon (significance denoted with a *). (P<0.05).

Other clinical signs were recorded as present or absent. Hamsters exhibited rough hair coat, orbital tightening, and hunched posture. Males displayed these more frequently than females in all groups, with six out of eight groups showing a significant sex difference (Fig. 1B). When observed over 7 dpi, males infected with Delta and Omicron had clinical signs less frequently than males infected with A.2.5, A.3, Beta, and Gamma. The Omicron-infected males also had less frequent clinical signs than Epsilon-infected males. Female hamsters exhibited no significant differences in clinical signs between variant groups.

#### Histopathology

Histopathological changes were evaluated to monitor respiratory disease progression 2, 4, and 7 dpi. Changes within the lungs were largely consistent across variants, with the majority of lesion variability observed between different dpi. At 2 dpi, lung lesions consisted of small to moderate amounts of intraluminal bronchial and bronchiolar inflammatory and epithelial cellular debris, intra-alveolar macrophages, neutrophils, necrotic cellular debris, fibrinous exudate, and hypertrophied vascular endothelial cells (Fig. 2A,B). In most variants, there was mild suppurative bronchitis and bronchiolitis, except for A.2.5 and Epsilon where these features were only present in one or two animals, respectively. Other variable changes across groups include bronchial epithelial hyperplasia, alveolar hemorrhage and edema, and perivascular lymphocytic aggregates. Epithelial syncytia were rare.

**FIGURE 2.**
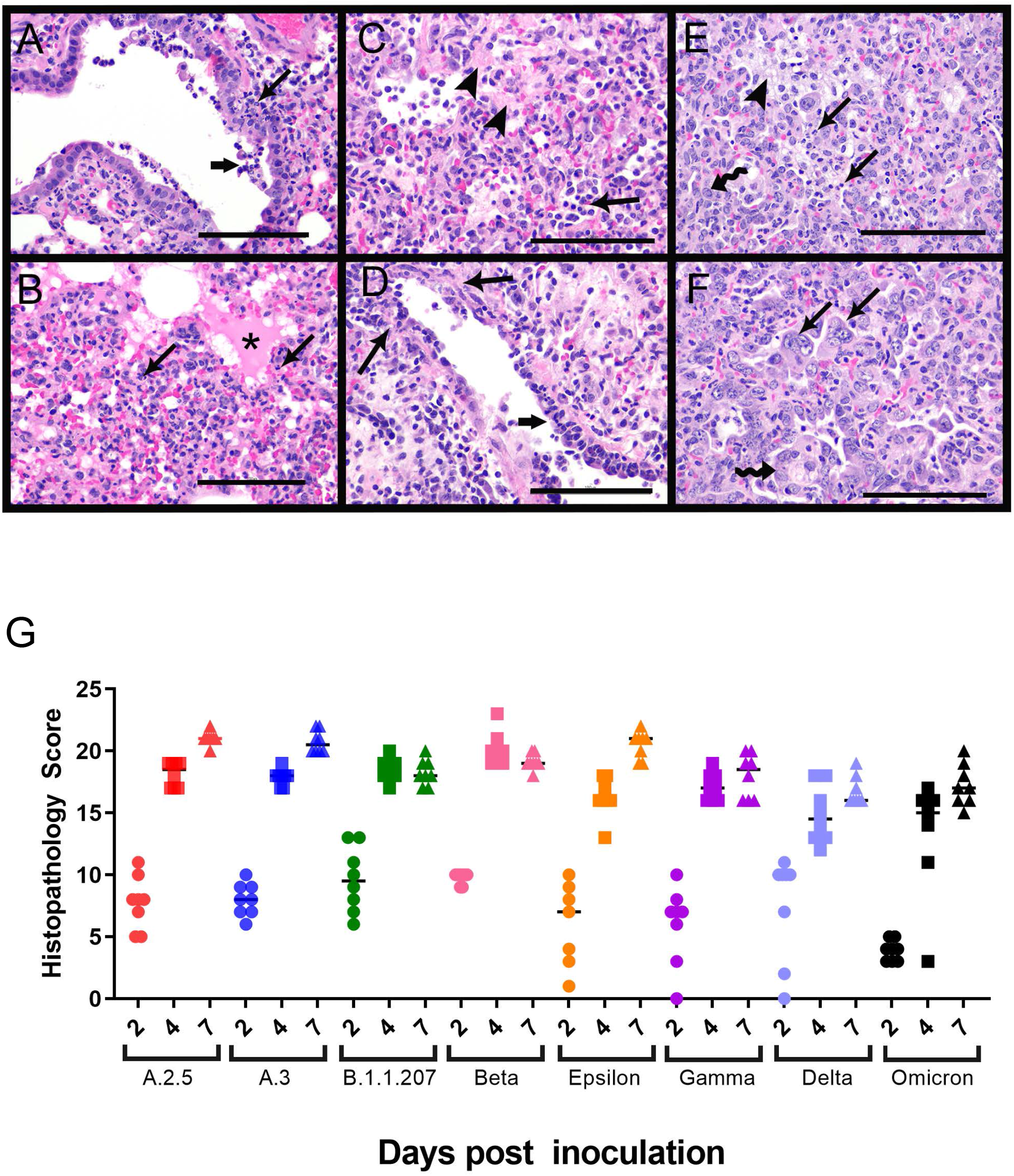
Histopathology of the lungs following naïve infection. Histopathologic changes of the lungs were consistent between variants at each time point. Representative images from each time point from different variants are shown. 2 dpi (A, B). (A) Suppurative bronchitis and bronchiolitis (long arrow) and bronchial epithelial cell necrosis along with inflammatory cell and necrotic cellular debris within the airways (short arrow) as observed in the Epsilon variant. (B) Alveolar edema (*) and inflammatory cells (arrows) including large numbers of neutrophils and macrophages as observed in Gamma variant. 4 dpi (C,D). (C) Organizing fibrin in the alveoli (arrowhead) and numerous inflammatory cells including neutrophils and macrophages (arrow) as observed in the B.1.1.207 variant. (D) Vasculitis/vascular endothelialitis characterized by loss of vessel wall integrity and eosinophilic proteinaceous material within the vessel wall (long arrows) and subendothelial mononuclear cells and neutrophils with endothelial cell swelling and necrosis (short arrow) as observed in the B.1.1.207 variant. 7 dpi (E,F). (E) Organizing fibrin within the alveoli (arrowhead), degenerate inflammatory cells within alveoli (arrows) and type II pneumocyte hyperplasia (squiggle arrow) as observed in the A.2.5 variant. (F) Multinucleated sloughed epithelial cells (arrows), type II pneumocyte hyperplasia (squiggle arrow), and intra-alveolar inflammatory cells as observed in the A.3 variant. Scale bars represent 100 μm. (G) Histopathology scores of naïve infection groups. Please see supplemental figure 2 for complete list of p values from multiple comparisons and one-way ANOVA. Each variant exhibited a similar trend with respect to scores at 2, 4, and 7dpi. Across time points, Omicron-infected animals had lower histopathology scores than A.3, B.1.1.207, and Beta 2dpi, all variants except Epsilon and Delta at 4 dpi, and all variants except B.1.1.207, Gamma, and Delta at 7dpi. Delta-infected animals had lower histopathology scores than B.1.1.207 and Beta at 4dpi and all variants except B.1.1.207, Gamma, and Omicron at 7dpi.

At 4 dpi, alveolar and perivascular edema was still appreciable and similar in severity to the edema seen at 2 dpi. However, the perivascular lymphocytic aggregates, intra-alveolar macrophages, neutrophils, necrotic cellular debris, and organized fibrinous exudate were more severe at 4 dpi than at 2 dpi. Additional lesions at 4 dpi included atypical type II pneumocyte hyperplasia and vasculitis (Fig. 2C,D). Vasculitis, when observed, was usually present in medium-sized arteries and veins and was characterized by loss of vessel wall integrity due to transmural effacement by inflammatory cells and necrosis, eosinophilic proteinaceous material within the vessel wall, or subendothelial mononuclear cells and neutrophils with endothelial cell swelling and necrosis. Similar lesions in human patients are described as endothelialitis (9), which has also been used in literature on hamster models (10–13). Suppurative bronchitis and bronchiolitis were variably present across groups at 4 dpi. Most animals infected with A.2.5, A.3, Epsilon, Gamma, and Delta demonstrated this lesion. However, fewer animals infected with Omicron and Beta variants, and no animals infected B.1.1.207 had this lesion.

At 7 dpi, the most consistent and striking lesion was areas of robust, atypical type II pneumocyte hyperplasia (Fig. 2E,F), which was seen in all animals and was usually present in at least 50% of tissue sections. Cells were up to 40 μm in diameter, cuboidal to polygonal, with round to oval nuclei up to 15μm in diameter that contained finely stippled chromatin and 1-3 prominent nucleoli. These cells had variable amounts of cytoplasm and occasionally were bi- or tri-nucleate, with numerous mitotic figures. Within the alveolar spaces and expanding the alveolar septa were large numbers of macrophages, with fewer neutrophils and lymphocytes, as well as fibrinous eosinophilic exudate. There was also bronchial and bronchiolar luminal necrotic cellular debris bronchial epithelial hyperplasia, perivascular lymphocytic aggregates, and vascular endothelial hypertrophy.

Most variants exhibited a similar histopathological trend, with scores increasing over time and no significant difference between sex appreciated (Fig. 2G). An exception were animals infected with B.1.1.207 and Beta variants, which had higher scores at 4 dpi than 7 dpi. These animals also had the highest 4 dpi scores across all variant groups. The lower scores at 7 dpi were due to lower percentage of tissue affected, smaller clusters of perivascular lymphocytes, and fewer animals with multinucleated or atypical bronchial epithelial cells. Omicron-infected animals had statistically lower histopathology scores than A.3, B.1.1.207, and Beta at 2 dpi, all variants except Epsilon and Delta at 4 dpi, and all variants except B.1.1.207, Gamma, and Delta at 7 dpi. Delta-infected animals had lower histopathology scores than B.1.1.207 and Beta at 4 dpi, and all variants except B.1.1.207, Gamma, and Omicron at 7 dpi. Despite overall lower histopathology scores, Omicron-infected animals demonstrated similar lesions with other variants (Supplemental Fig. 1).

#### Viral titers

Lung, tracheal, and nasal turbinate titers were measured to evaluate viral tropism in respiratory tissues over time. Overall, the highest viral titers were observed at 2 dpi in all tissue types, with lungs and nasal turbinates having higher titers than the trachea (Fig. 3A-C). All variant groups displayed a quick decrease in infectious virus load at 4 dpi, while no infectious virus was detected in the lung or tracheal samples at 7 dpi (Fig. 3A-C).

**FIGURE 3.**
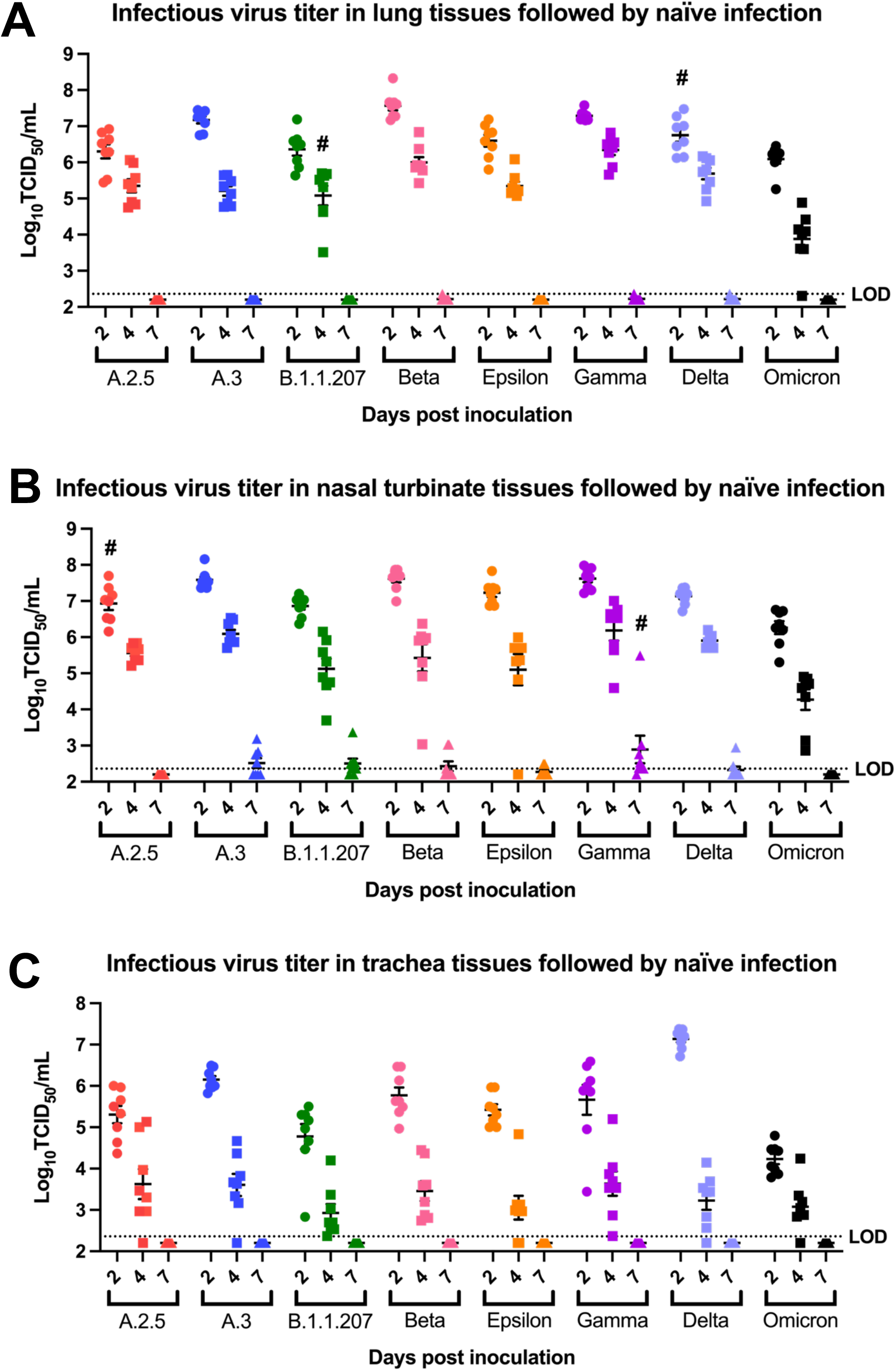
Viral titers in respiratory tissues following naïve infection. (A) Mean infectious viral titers in lung tissues at 2 dpi, 4 dpi, and 7 dpi after naïve infection (n=8). Significant differences in tissue viral titers were observed between some variants at 2 dpi and 4 dpi. Notably Beta and Gamma had higher titers and Omicron had lower titers at both 2 dpi and 4 dpi. No significant difference was observed at 7 dpi. Groups that displayed sex differences in viral titers were indicated with # sign with males being higher than females. (B) Mean infectious viral titers in trachea tissues at 2 dpi, 4 dpi, and 7 dpi after naïve infection (n=8). Delta had the highest titers in all variants tested, and Omicron had lower titers than all other variants at 2 dpi. No significant difference in tracheal titers were observed between variants at 4 dpi, 7 dpi, or between sex at all timepoints. (C) Mean infectious viral titers in nasal turbinate tissues at 2 dpi, 4 dpi, and 7 dpi after naïve infection (n=8). Few significant differences in viral titers were observed between variants, apart from Omicron having lower titers at 2 dpi and 4 dpi. No significant difference was observed at 7 dpi. Groups that displayed sex differences in viral titers were indicated with # sign with males being higher than females. Viral titer was measured by TCID_50_ assay with limit of detection at 2.36 log_10_TCID_50_/mL. One-way ANOVA was done on all three tissue types with Turkey’s multiple comparison tests. (p < 0.05) Please see supplemental table 4 for complete list of p-values for multiple comparison and one-way ANOVA.

Significant differences in mean viral titers were observed between variants for all three tissue types, with Omicron-infected animals displayed consistently lower tissue titers than other groups. In lung tissues, the most prominent difference was noted at 4 dpi, with the Omicron group having significantly lower lung titers than all other groups, while Beta and Gamma groups had significantly higher titers than all groups except Delta (Fig 3A). At 2 dpi, this trend was still appreciable and Omicron-infected animals had significantly lower titers than other variants except A.2.5, B.1.1.207, and Epsilon, which were not significantly different (Fig. 3A, Supplemental table 4). Beta-infected animals had the highest overall lung titer and, along with Gamma-infected animals, had significantly higher titers than those infected with A.2.5, B.1.1.207, Epsilon, and Omicron. Delta-infected animals had comparable lung titers to those infected with A.2.5, A.3, B.1.1.207, Epsilon, and Gamma, while being significantly lower than Beta- and higher than Omicron-infected animals. Delta was the only variant with sex differences in lung titers at 2 dpi, with males having significantly higher mean titers than females (Fig. 3A). At 4 dpi, the only sex difference appreciated was B.1.1.207-infected males having significantly higher titers than females (Fig. 3A). Altogether, infectious viral titers in lung tissues were higher in Beta-infected animals and lower in Omicron-infected animals. These results correlate well with histopathology findings, suggesting an early infection followed by a reparative phase (Fig. 2G).

In nasal turbinates at 2 dpi, Omicron-infected animals had significantly lower viral titers than those infected with A.3, Beta, and Gamma (Fig. 3B). Although A.3, Beta, and Gamma groups trended towards having higher nasal turbinate titers at this timepoint, statistical tests suggested no significance. A.2.5-infected animals displayed sex differences in nasal turbinate titers at 2 dpi, with males having higher viral titer than females (Fig. 3B). At 4 dpi, Omicron-infected animals continued to have the significantly lowest nasal turbinate viral titer. Gamma-infected animals had the highest, with significantly higher titers than B.1.1.207, Epsilon, and Omicron-infected animals. At 7 dpi, low nasal turbinate titers were detectable for a few animals from multiple variant groups, with the exception for Gamma which had detectable virus in 7 out of 8 hamsters’ nasal turbinates (Fig. 3B). Within the Gamma-infected animals at this timepoint, males demonstrated a significantly higher nasal turbinate titer than females.

At 2 dpi, Omicron-infected animals had the significantly lowest tracheal viral titer while Delta-infected animals exhibited the highest (Fig. 3C). Tracheal tissue viral titers at 4 dpi had more variability between variants than other tissue types, but these differences were not statistically significant. There were no sex differences in tracheal titers at all timepoints.

### Reinfection

#### Clinical outcomes

To investigate if previous SARS-CoV-2 infection ameliorates the frequency of clinical signs observed after reinfection, hamsters were inoculated with variant A and then a different variant 28 days later. After the first infection, hamsters lost up to 24% BW over the first 6 dpi, and then began gaining weight around 7 dpi. They continued to gain BW until reinfection, after which their weights plateaued or slightly decreased immediately following inoculation, before increasing again typically around 2 or 3 dpi. Supplemental figure 2A-H compares these trends in percent BW change after reinfection to that seen with naïve infected animals for each variant. Figure 4A displays the difference in BW change for each hamster between their initial and reinfection, with significantly less weight loss seen after reinfection, except for animals infected with variant A.3 at 28 dpi which showed no significant difference. The non-significance for the A.3 reinfected hamsters may be attributed to one hamster that exhibited a weight loss pattern similar to a naïve infected animal. The mock inoculated group lost significantly more BW following infection at 28 dpi than other groups. The mean percent BW change after reinfection was also significantly different from naïve animals infected with the same variant, with naïve animals losing more BW (Fig. 4B). There were no sex differences in BW change in these groups following reinfection.

**FIGURE 4.**
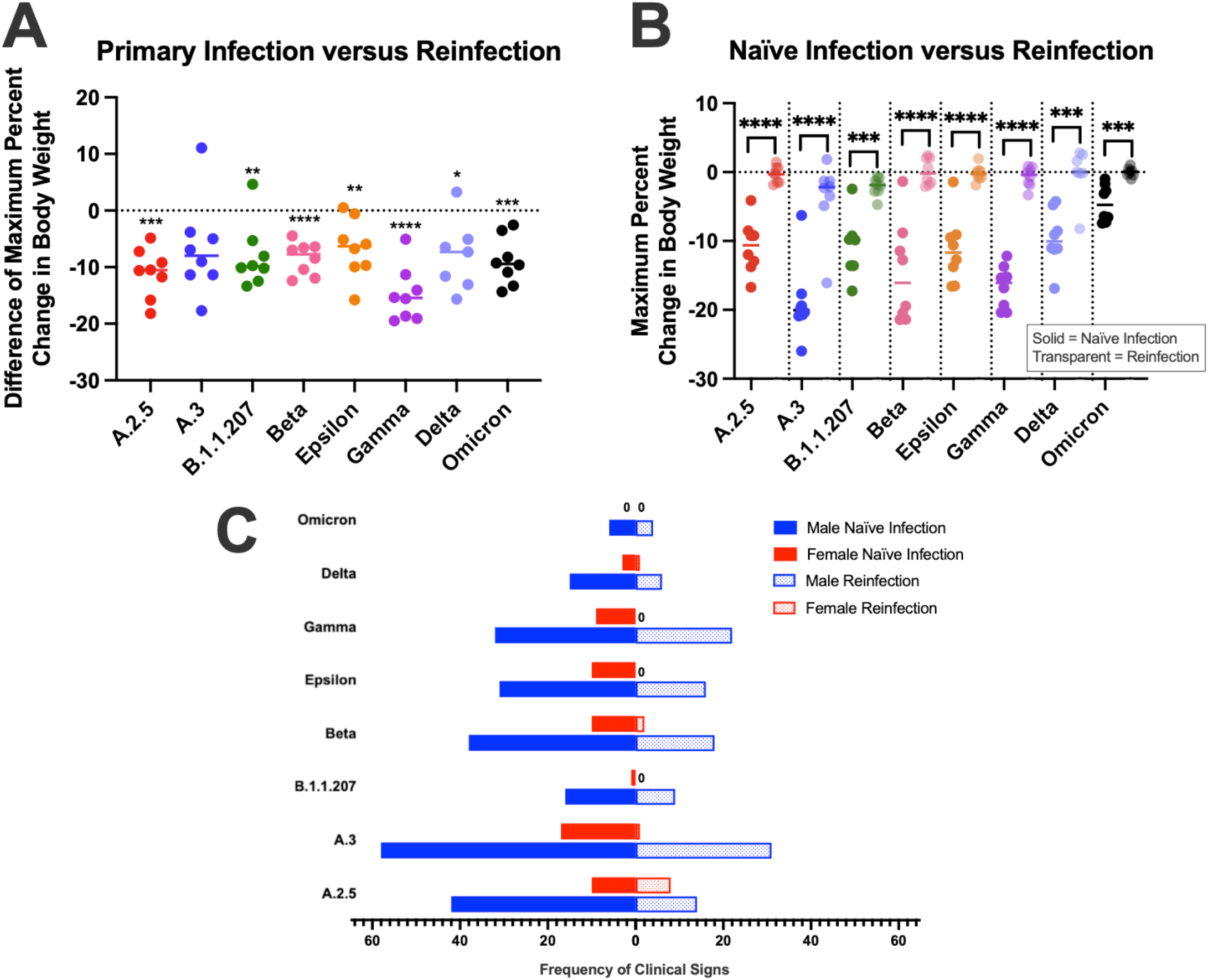
Clinical signs following reinfection. (A-B) The maximum percent body weight change over 7 dpi. (A) The difference in each hamster’s weight change between primary infection with variant A and reinfection with another variant was compared using paired t-test, with almost all animals showing significantly less weight loss after reinfection. (B) When compared using two sample t-tests, naïve hamsters infected with a variant lost significantly more weight than their counterparts that were reinfected with a variant after previous infection with variant A. (C) The frequency of clinical signs (hunched posture, orbital tightening, and rough hair coat) is shown for naïve male and female hamsters 7 dpi as well as male and female hamsters 7 dpi reinfection. Clinical signs were observed less in reinfected male hamsters versus naïve male hamsters except for Omicron and Delta infected animals. A.3 and Gamma reinfected females had significantly less clinical signs than their naïve counterparts. (P <0.05).

**FIGURE 5.**
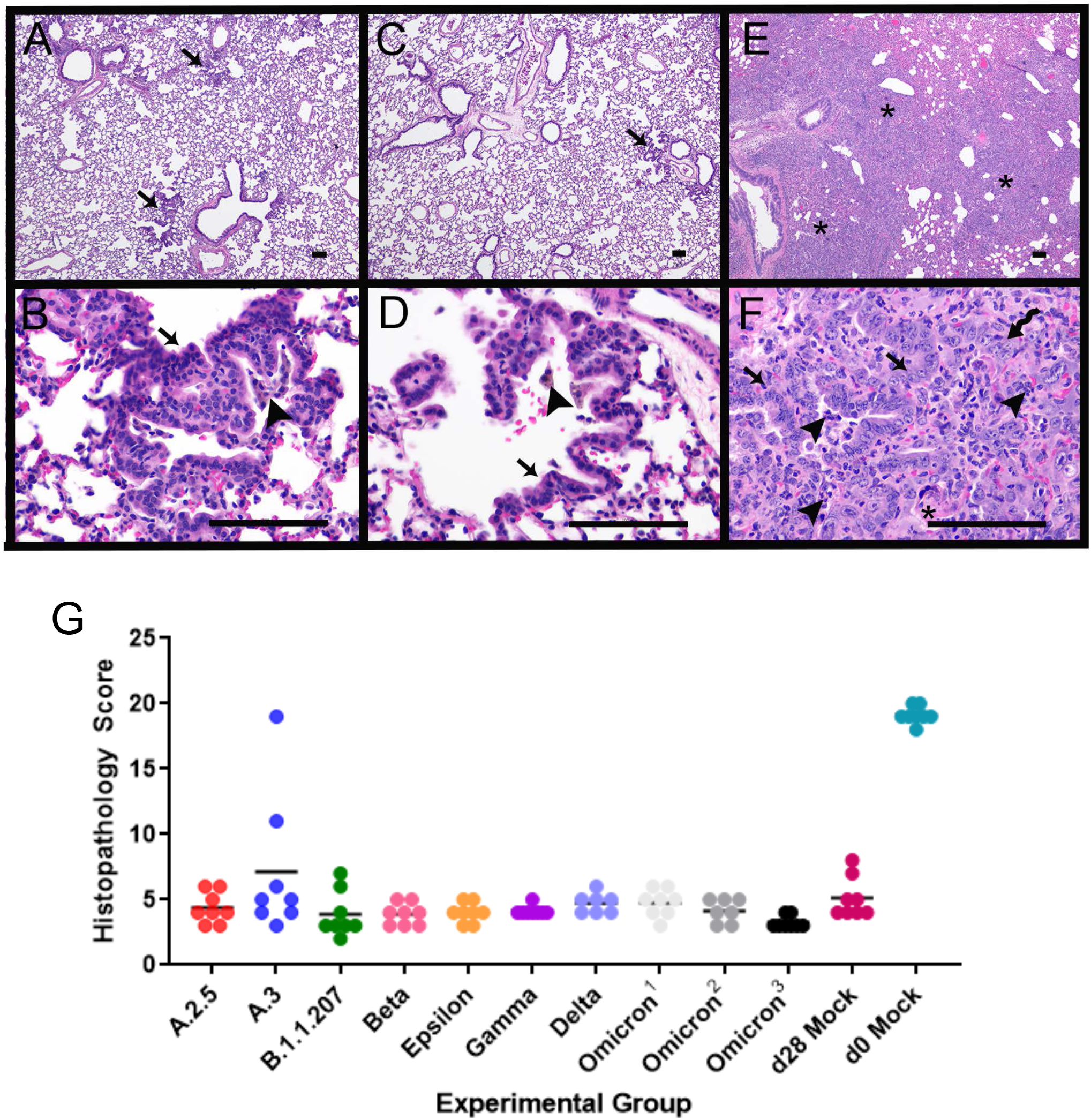
Histopathology of the lungs following reinfection. Representative images from reinfection with Delta variant (A,B). (A) Low magnification showing largely unremarkable lung with few foci of residual type II pneumocyte hyperplasia (arrows). (B) Higher magnification showing type II pneumocyte hyperplasia (arrow) and clusters of intra-alveolar clusters of pigmented macrophages (arrowhead). Representative images from mock reinfection (C,D). (C) Low magnification showing largely unremarkable lung with foci of residual type II pneumocyte hyperplasia (arrow). (D) Higher magnification view of type II pneumocyte hyperplasia (arrow) and intra-alveolar pigmented macrophage (arrowhead). Representative images from day 0 mock infection with variant reinfection (E, F). E. Low magnification view indicating widespread consolidation of the alveolar spaces (*). (F) Higher magnification showing type II pneumocyte hyperplasia (arrows), intraalveolar inflammatory cells including neutrophils and macrophages (arrowheads), organizing fibrin within alveoli (*), and multinucleated epithelial cells (squiggle arrow). All scale bars represent 100 μm. (G) Histopathology scores of reinfection groups. Animals were intranasally infected with A variant at day 0, infected with an additional variant at 28 dpi, and euthanized at 35 dpi. ^1.^ Initial infection with Delta, reinfection with Omicron. ^2.^ Initial infection with Mu, reinfection with Omicron. ^3.^ Initial infection with A, reinfection with Omicron. D28 mock: animals were infected with A variant at day 0, and mock-inoculated with media only at 28 dpi. D0 mock: animals were mock-inoculated with media only at day 0, and inoculated with A variant at 28 dpi. Please see supplemental table 3 for complete list of p values from multiple comparisons and one-way ANOVA. Secondary variant infection at 28 dpi produced similar histopathology scores. D0 mock histopathology scores were statistically significantly higher (p < 0.0001, multiple comparisons one-way ANOVA) than all other groups. Variant A.3 had 2 animals with high histopathology scores, but scores were otherwise similar to all other groups.

The presence and frequency of clinical signs were recorded over 7 dpi for both infections as described above. Male hamsters displayed more frequent clinical signs (rough hair coats, orbital tightening, and hunched posture) than females after both initial and reinfections (Fig. 4C). Significantly fewer clinical signs were observed after reinfection for males infected with A.2.5, B.1.1.207, Delta, and Omicron, and females infected with Gamma. Almost all male groups displayed fewer clinical signs following reinfection when compared to naïve males infected with the same variant, except for B.1.1.207, Delta, and Omicron that had a low frequency of clinical signs for both infections (Fig. 4C). Females previously infected with A.3 and Gamma also had significantly fewer clinical signs compared to their naïvely infected counterparts.

To evaluate if the initial variant influences reinfection outcomes, separate cohorts of animals were reinfected with Omicron 28 days after initial infection with Delta or Mu. These hamsters exhibited similar weight change trends to hamsters infected with variant A before Omicron reinfection (Supplemental fig. 3A,B). There were no significant differences in weight change between reinfected hamsters based on the variant they were initially infected with; however, all three groups of hamsters reinfected with Omicron had significantly less BW change compared to naïve Omicron-infected hamsters. The frequency of clinical signs between reinfection groups was not significantly different.

#### Histopathology

Histopathology was performed on lung tissues at 7 dpi following reinfection (35 dpi initial infection, thus 28 days apart). Animals had largely unaffected lungs 7 days after reinfection. In all variant groups, there were clusters of regular, cuboidal type II pneumocyte hyperplasia, admixed with pigmented macrophages. Less consistent features across variants include alveolar hemorrhage and edema and perivascular lymphocytic aggregates.

In animals mock-inoculated at day 28, there were clusters of type II pneumocyte hyperplasia with aggregates of pigmented macrophages, similar to hamsters that underwent two infections. In animals that were mock inoculated at day 0, and inoculated at day 28 with variant A, there was robust atypical type II pneumocyte hyperplasia as described previously in the 7 dpi pathogenesis groups, along with abundant intra-alveolar and intraseptal macrophages. The day 0 mock-inoculated animals had statistically higher histopathology scores than all other groups. Animals infected with variant A.3 had statistically higher histopathology scores than the other groups, but this was largely attributed to one animal with widespread atypical type II pneumocyte hyperplasia and alveolar infiltrates as described for 7 dpi pathogenesis animals.

Tissue viral titer analyses were not performed for these groups as animals were euthanized at 7 days after reinfection, and low tissue viral load at 7 dpi from this study and others have been reported (6, 14).

## Discussion

SARS-CoV-2 variants have emerged with mutations affecting viral ACE2 binding affinity, replication, transmission, antibody recognition and clinical disease (1, 15). Validating animal models that accurately demonstrate disease caused by these variants is important for studying variant pathogenesis and assessing efficacy of therapies and vaccines. Here, Syrian hamsters infected with SARS-CoV-2 variants displayed clinical outcomes, lung pathology, and tissue viral titers as previously described (16–18), with some differences observed between variants and between sexes within variant groups.

BW loss was a reliable clinical indicator of SARS-CoV-2 infection in this study. It was observed across variants regardless of sex, with naïve hamsters losing weight over the first 6 dpi before beginning to recover on 7 dpi. This is consistent with previous studies reporting around 10-14% BW loss (7, 13, 14, 16–20) and similar patterns of recovery (6, 18). While all groups followed this pattern, some variant groups exhibited significant BW differences. A.3-infected animals demonstrated the most significant BW loss, particularly in males. This was an unexpected outcome as A.3 is genetically similar to the originally circulating virus with minimal mutations, and the groups that exhibited relatively less weight loss were infected with variants possessing multiple mutations known to affect viral fusogenicity and infectivity (1, 15, 21, 22). Previous studies with a similar lineage did not report this severity of BW loss despite similar lung pathology (6, 14).We postulate that these differences could be due to variations in inoculation procedures or viral preparation. Despite the increased severity of clinical disease, A.3-infected animals had similar lung histopathology scores and tissue viral titers to most other groups. While this variant is no longer dominantly circulating among humans, our findings are useful as they highlight potential differences in SARS-CoV-2 infection between hamsters and humans.

The groups with the least BW loss were infected with Omicron, which is consistent with previous reports of Omicron clinical presentations in hamsters and human patients (23–28). In one study comparing Delta and Omicron infection in Syrian hamsters, Omicron caused less BW loss and lower viral loads in throat swabs and nasal wash samples (29). Due to its dominance in the population, Omicron is a priority for therapeutic and preventative research, particularly as it continues to evolve into multiple sub-lineages. In this study, we evaluated the early sub-lineage BA.1, referred to as Omicron throughout this paper.

The etiology of BW loss in infected hamsters is not entirely understood but it likely recapitulates the condition in human patients. Weight loss and clinical cachexia in people can be attributed to factors like loss of appetite and taste, anosmia, fever and inflammation, and metabolic imbalances (30, 31). While we did not evaluate appetite, taste, or smell, anosmia in infected hamsters has been reported to occur at 2 to 5 dpi (32, 33), which correlated with the period of maximal weight change (6, 18, 32, 33). Meanwhile, there is limited description on SAR-CoV-2 effects on hamster core body temperatures, but one study reported no changes following infection with an early variant (34). We postulate that the weight loss in hamsters is associated with inappetence, which could be a result of anosmia as correlated with the histological damage within the olfactory epithelium (32). Additional studies examining the relationship of weight loss with these factors may elucidate its etiology.

The other clinical signs observed across variant groups were consistent with previous studies (16, 18); however, respiratory signs like rapid breathing have also been reported (17). As with our study, the majority of the literature describes minimal (6) to no (16) respiratory signs(13, 18). Such clinical severity differences could be attributed to factors like viral inoculation dose, with higher doses resulting in increased morbidity (17).

The influence of sex on clinical sign frequency in our animals mirrors the increased COVID-19 morbidity documented in male human patients (35–39). Male hamsters displayed more frequent clinical signs than females, which is also consistent with previous reports of increased morbidity in male hamsters (14, 38, 40). Moreover, inter-variant differences in clinical sign frequencies were observed in male hamsters, suggesting they may be more sensitive to phenotypic effects of SARS-CoV-2 infection than females, both in the context of the initial infection and after reinfection. Of note, despite differences in the overall clinical sign frequencies, there were no sex differences associated with BW change up to 7 dpi for all groups, which is consistent with previous reports (14, 17, 38)<u>; h</u>owever, some studies of longer duration noted that male hamsters regained less weight than females from 8 to 28 dpi (14, 38). Sex differences emphasize the importance of accounting for sex in SARS-CoV-2 research.

Overall, histopathological features were similar among all variants and consistent with previous literature (6, 7, 10, 13, 20, 41–43). Inflammatory lesions present at 2 dpi supported acute damage, which then progressed in severity by 4 dpi. In some animals, evidence of repair such as type II pneumocyte hyperplasia was already present by 4 dpi, but there was no obvious correlation to sex or variant with these repairs. At 7 dpi, lesions were consistent with further progression into the reparative phase, as characterized by extensive type II pneumocyte hyperplasia. In most variants, lesions were more extensive at 7 dpi; however, there were more features present at 4 dpi. As a result, two variants (B.1.1.207 and Beta) had higher histopathology scores at 4 dpi than 7 dpi. The histopathologic progression is consistent with the weight trends observed, with an immediate response to infection and gradual recovery towards 7 dpi. In people, respiratory lesions are primarily characterized by imaging such as computed tomography (CT) (44, 45). The sensitivity of CT for diagnosing infection in humans increases significantly when symptom duration is longer than 48 hours, after which increased lung consolidation and ground glass opacities are observed (45). CT has been used to evaluate SARS-CoV-2 in hamsters with similar findings to human patients (7), and offers a viable option for evaluating lung pathology over time in this model (14, 18).

Omicron-infected animals had lower overall histopathology scores compared to most variants despite similar lesions observed (Supplemental Fig. 1), likely due to lower percentages of affected tissue, fewer perivascular lymphocytes, and fewer atypical or multinucleated bronchial epithelial cells. Our findings differed slightly from previous reports that used lower inoculation doses, where hamsters demonstrated milder pneumonia when infected with Omicron compared to the Delta variant (23, 26). These studies found that Omicron-infected animals had milder features of pneumonia, including multiple small foci of inflammatory cells in the alveoli and peribronchial areas only observed at 6 dpi, with no changes noted at 3 dpi (26), or decreased areas of type II pneumocyte hyperplasia at 5 and 7 dpi (23).

All variant groups displayed a quick decrease in infectious virus load in respiratory tissues from 2 dpi to 4 dpi, and by 7 dpi, there were low to no detectable infectious viral particles. This pattern is similar to previous reports for hamsters infected with a Clade A variant (6, 14, 17). The viral titer levels at 7 dpi coincide with the peak BW loss at 6 dpi and then recovery observed in this study and others (6). However, higher inoculation doses could result to consistently higher viral titer loads at 2 and 4 dpi, as previously reported (Chan et al. 2020). In our study, the high titer levels in the nasal turbinates and lungs may be attributed to the nasal cavity inoculation and more effective viral replication at 37°C in the lower respiratory track, respectively.

There was a reduction in clinical phenotype and lung pathology upon SARS-CoV-2 reinfection. Previously infected hamsters had significantly less BW loss than their naïvely infected counterparts, and the majority of groups had less frequent other clinical signs. Additionally, in 79/80 animals that were reinfected, lesions at 35 dpi (i.e., 7 days post-reinfection) were identical to those of the control group that received only one infection with variant A on day 0 and were mock inoculated at 28 dpi. These changes included residual type II pneumocyte hyperplasia and small foci of perivascular and intra-alveolar inflammatory cells, which were consistent with chronic, rather than acute, change. In contrast, animals mock-inoculated at day 0 and inoculated at 28 dpi displayed changes more like those observed in 7 dpi animals. One animal in the reinfection cohort demonstrated findings similar to those found in animals euthanized at 7 dpi. Serology performed at 28 dpi on this individual revealed no neutralizing antibodies (data not shown) indicating that there was likely a failure of primary inoculation. Protective antibody responses of variable durations are described in human cases, both from previous infection and vaccines (46–48). Overall, our findings indicate that some degree of protection against reinfection is generated during primary infection.

A.3 and Omicron-infected groups demonstrated clear phenotypic differences from other groups, supporting the recommendation that all variants should be characterized in the hamster model. Despite differences in reported transmissibility, viral replication, and clinical outcomes in humans and *ex vivo* studies(1, 2), the other variants analyzed were phenotypically similar. This suggests that although there are benefits as an animal model overall, the hamster model may not be specific enough to differentiate minor differences between certain variants. Despite this, preliminary studies on emerging variants using the model are still valuable for characterization before further investigation is conducted.

While we evaluated the effect of sex on different variants, there are other factors that can influence SARS-CoV-2 infection. Age has been shown to significantly affect disease outcomes in people (49–51) and in hamsters (10, 52). An additional limitation of our study is its duration; longer studies are needed to investigate effects of long COVID and the longevity of the protection from reinfection. Furthermore, evaluation of the hamster antibody response to different variants is important, as hamsters continue to be used for vaccine and mAb treatment research (53–55).

Emergence of new variants creates a continued need for SARS-CoV-2 research. Our study findings indicate that Syrian hamsters provide a reliable and consistent animal model for studying SARS-CoV-2 variant infections and reinfections, emphasizing its significance for characterizing disease and investigating effective treatments and vaccines.

## Materials and Methods

### Virus Preparation

Virus preparation was performed as previously described by Mulka *et al*. (2021). Briefly,.Vero-E6-TMPRSS2 cells (from the Japan Institute of Infectious Diseases) were cultured in DMEM supplemented with 10% fetal bovine serum (FBS), 1mM glutamine, 1mM sodium pyruvate, and penicillin (100U/ml) and streptomycin (100mg/ml). SARS-CoV-2 viruses from patient nasal swab samples collected at Johns Hopkins Hospital were used to infect Vero-E6-TMPRSS2 cells to generate virus stocks. Infection was done at a multiplicity of infection (MOI) of 0.01 TCID_50_. Infected cell supernatant was collected when 75% of cells had observable cytopathic effect (CPE) around 96 hours post infection. The supernatant was centrifuged at 400x g for 10 min and then aliquoted into 500uL and stored at −70°C. Infectious titer of virus stock was measured by TCID_50_ assay. Virus stocks were 10-fold serially diluted in DMEM supplemented with 2.5% FBS, 1mM glutamine, 1mM sodium pyruvate, and penicillin (100U/ml) and streptomycin (100mg/ml). Stock dilutions were transferred in sextuplicate into the 96-well plates confluent with Vero-E6-TMPRSS2 cells, incubated at 37°C for 6 days, then fixed with 4% formaldehyde and stained with naphthol blue black solution for visualization. The infectious virus titers in TCID_50_/ml were determined by the Reed and Muench method.

### Animals

Male and female Syrian hamsters (6-8 wk old, Envigo, Haslett, MI) were singly housed in negative-pressure individually ventilated cages (PNC, Allentown, Allentown, NJ). Animals were provided with corncob bedding (Envigo, Madison, WI), nesting material (Enviro-dri, Shepherd Specialty Papers, Amherst, MA), standard rodent chow (2018SX, Teklad, Envigo, Madison, WI) and RO-DI water through an automated watering system (Edstrom, Avidity Science, Waterford, WI). All animal procedures were approved by the Johns Hopkins Institutional Animal Care and Use Committee and conducted in an AAALAC-accredited facility.

#### Inoculation, clinical evaluation, and euthanasia

Hamsters were sedated with ketamine/xylazine for inoculation with 10^5^ TCID_50_ of a SARS-COV-2 variant (Table 1) in 100ul DMEM (50 ul/naris) or mock-inoculated with 100ul DMEM. Animals were weighed and clinically evaluated by a blinded observer (JP). At euthanasia, hamsters were anesthetized using isoflurane for cardiac puncture and tissue harvest.

#### Naïve SARS-COV-2 Variant Infection

Hamsters (4/sex/endpoint/variant) were inoculated with a SARS-COV-2 variant. Daily observations and weighing were performed until euthanasia at of 2, 4, and 7 dpi.

#### Variant Reinfection

For primary infection, hamsters (4/sex/variant) were inoculated with 10^5^ TCID50 of SARS-CoV-2 130 USA-WA1/2020. Negative control animals (4/sex) were mock inoculated. 28 days after induction of primary infection, experimental groups were inoculated with 10^5^ TCID_50_ of a variant (Table 1) and the initially mock infected animals were inoculated with 10^5^ TCID_50_ of SARS-CoV-2 130 USA-WA1/2020. Periodic clinical assessment was performed until euthanasia at 35 dpi.

#### Delta versus Mu effect on Omicron infection

Hamsters (4/sex/variant) were inoculated with 10^5^ TCID_50_ of SARS-CoV-2/USA/MD-HP06587-PIDGNNWCBG/2021 or 10^5^ TCID50 of SARS-CoV-2/USA/MD-HP05660/2021. At 28 dpi, hamsters were inoculated again with 10^5^ TCID_50_ of SARS-CoV-2/USA/MD-HP20874-PIDUYWZOWA/2021. Clinical assessment and euthanasia were performed following the timepoints for the reinfection study above.

### Histopathologic Analyses

Histopathological analysis of lung tissues on H&E slides was performed blindly by a veterinary pathologist (KM) using a scoring system (Table S1)(12, 56, 57). Briefly, parameters evaluated for their presence or absence include: necrosis of bronchiolar epithelial cells (BEC), cellular debris in bronchi and bronchioles, cellular debris in alveoli, intra-alveolar fibrin, alveolar hemorrhage, alveolar edema, perivascular or interstitial edema, vasculitis, plump vascular endothelial cells, necrosuppurative bronchitis, hyperplasia of BEC, hyperplasia of type II alveolar epithelial cells (AEC), multinucleated or atypical BEC, and multinucleated or atypical AEC. Factors evaluated on a scale from 0-4 included percent of lung affected and perivascular lymphocytes. Intra-alveolar neutrophils and macrophages were evaluated on a scale from 0-3.

### Tissue Homogenization and Viral Titer Analysis

Animal tissue homogenization and infectious viral titration were done as previously described (Mulka *et al*. 2021, Dhakal *et al*. 2021). Briefly, tissue samples were transferred to Lysing Matrix D beads tubes on ice. DMEM supplemented with penicillin (100U/ml) and streptomycin (100mg/ml) was added to the tubes at 10% wt/vol ratio. The samples were loaded in a FastPrep-24 bench-top bead beating system (MPBio) and homogenized for 40 seconds at 6.0 m/s, followed by centrifugation for 5 min at 10,000x g at room temperature. Tubes were returned to ice, and supernatant was collected and stored at −70°C. Infectious virus titer in tissue homogenates was measure by TCID_50_ assay as described above.

### Statistical Analysis

Statistical analyses were performed using GraphPad Prism 9. Paired t-tests were used to evaluate differences in BW and clinical signs in reinfection groups. One-sample t-tests were used to evaluate sex differences within groups and clinical outcome differences between naïve and reinfection. One-way analysis of variances (ANOVA) were used to evaluate differences between variant group BW, clinical signs, tissue titers, and histology scores. A significance of P <0.05 was used for all tests.

## Acknowledgements

This study was funded by the Centers of Excellence for Influenza Research and Response (NIAID N272201400007C) and the Richard Eliasberg Family Foundation. We also thank Jacqueline Brockhurst, Natalie Castell, Morgan Craney, Isabel Jimenez, Amanda Maxwell, Andrew Johansen, Lyle Nyberg, and Riley Richardson.

---

**SUPPLEMENTAL FIGURE 1.**
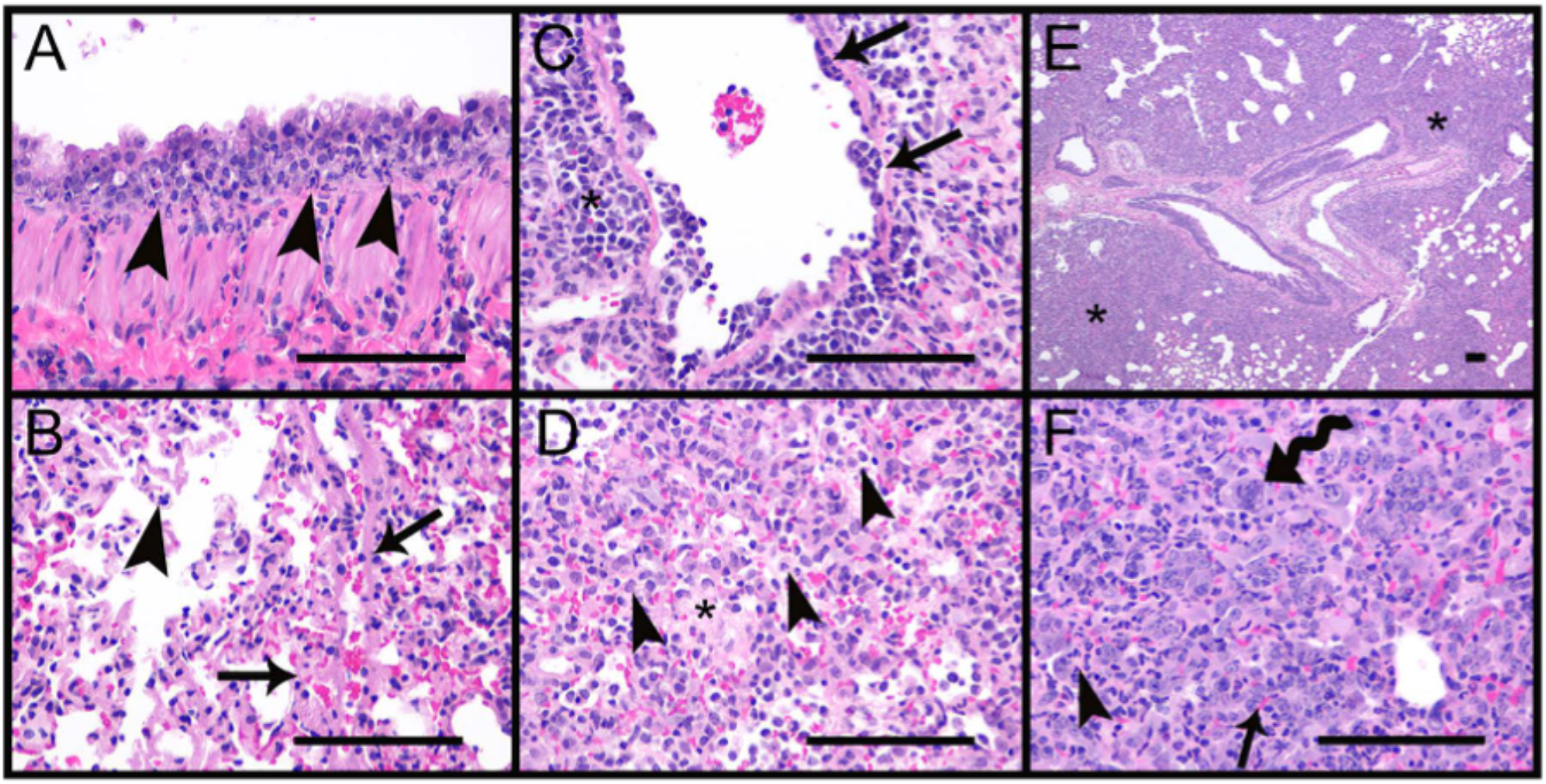
Histopathology of the lungs, Omicron variant. 2 dpi (A,B). (A) Suppurative bronchitis (arrowheads), with bronchial epithelial cell degeneration and necrosis. (B) Intra-alveolar organized fibrin (arrows), and increased inflammatory cells including neutrophils and macrophages (arrowhead) within the alveolar septa and spaces. 4 dpi (C,D). (C) Vasculitis/vascular endothelialitis characterized by subendothelial aggregates of neutrophils and mononuclear cells (arrows), as well as perivascular accumulation of inflammatory cells including neutrophils and macrophages (*). (D) Organized fibrin (*) within the alveoli along with increased numbers of inflammatory cells including neutrophils and macrophages (arrowheads). 7 dpi (E, F). (E) Low magnification indicating the extent of the consolidation (*) of the alveoli. (F) Higher magnification indicating type II pneumocyte hyperplasia (arrow), inflammatory cells including neutrophils (arrowhead), macrophages, and fewer lymphocytes, and multinucleated epithelial cells (squiggle arrow). Scale bar represents 100 μm in all images.

**SUPPLEMENTAL FIGURE 2.**
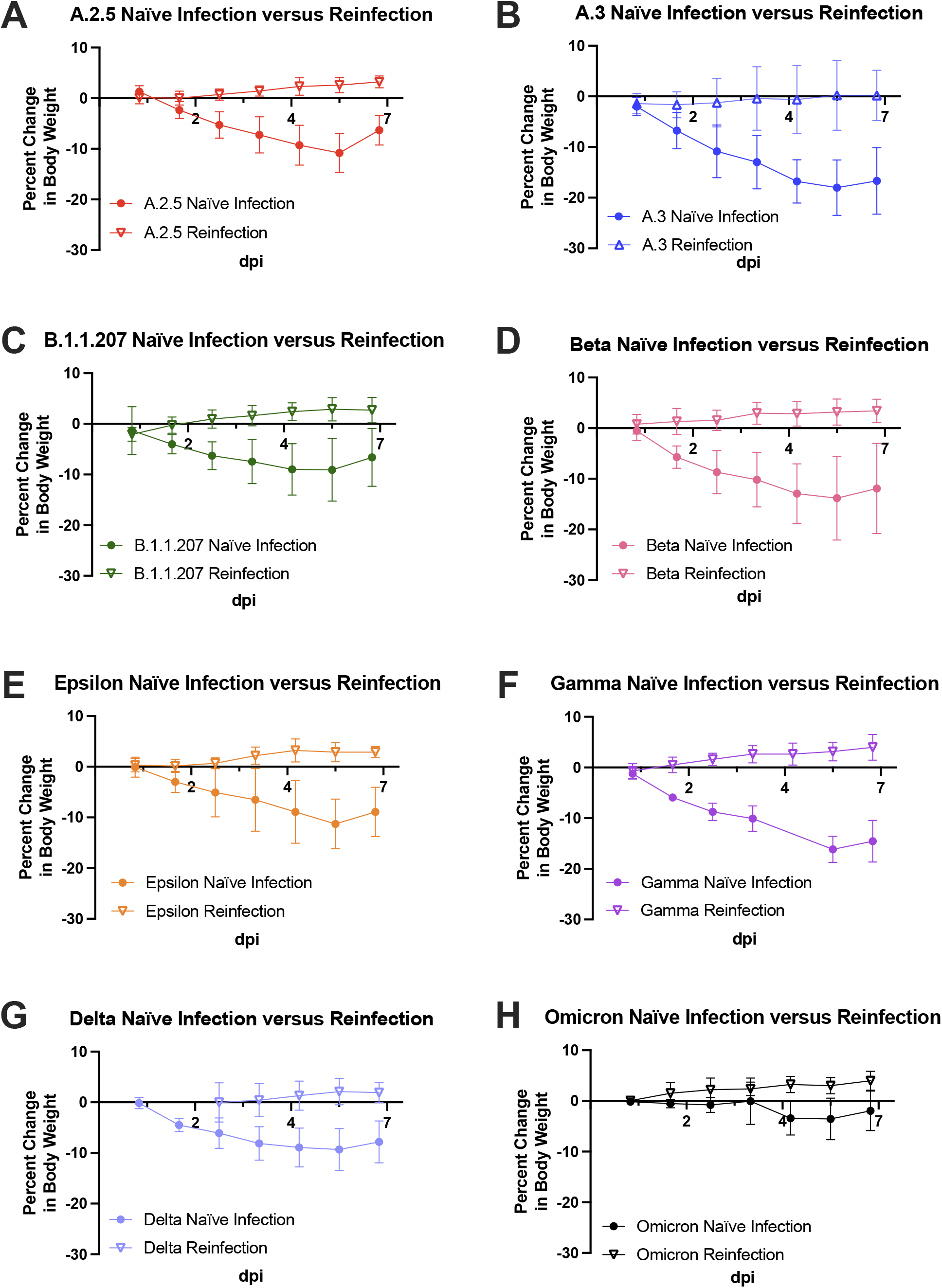
Percent weight change over 7 days following variant naïve infection and variant reinfection. (A-H) Naïve hamsters consistently lost weight after infection, while reinfected hamsters showed little to no initial weight loss followed by weight gain over 7 dpi.

**SUPPLEMENTAL FIGURE 3.**
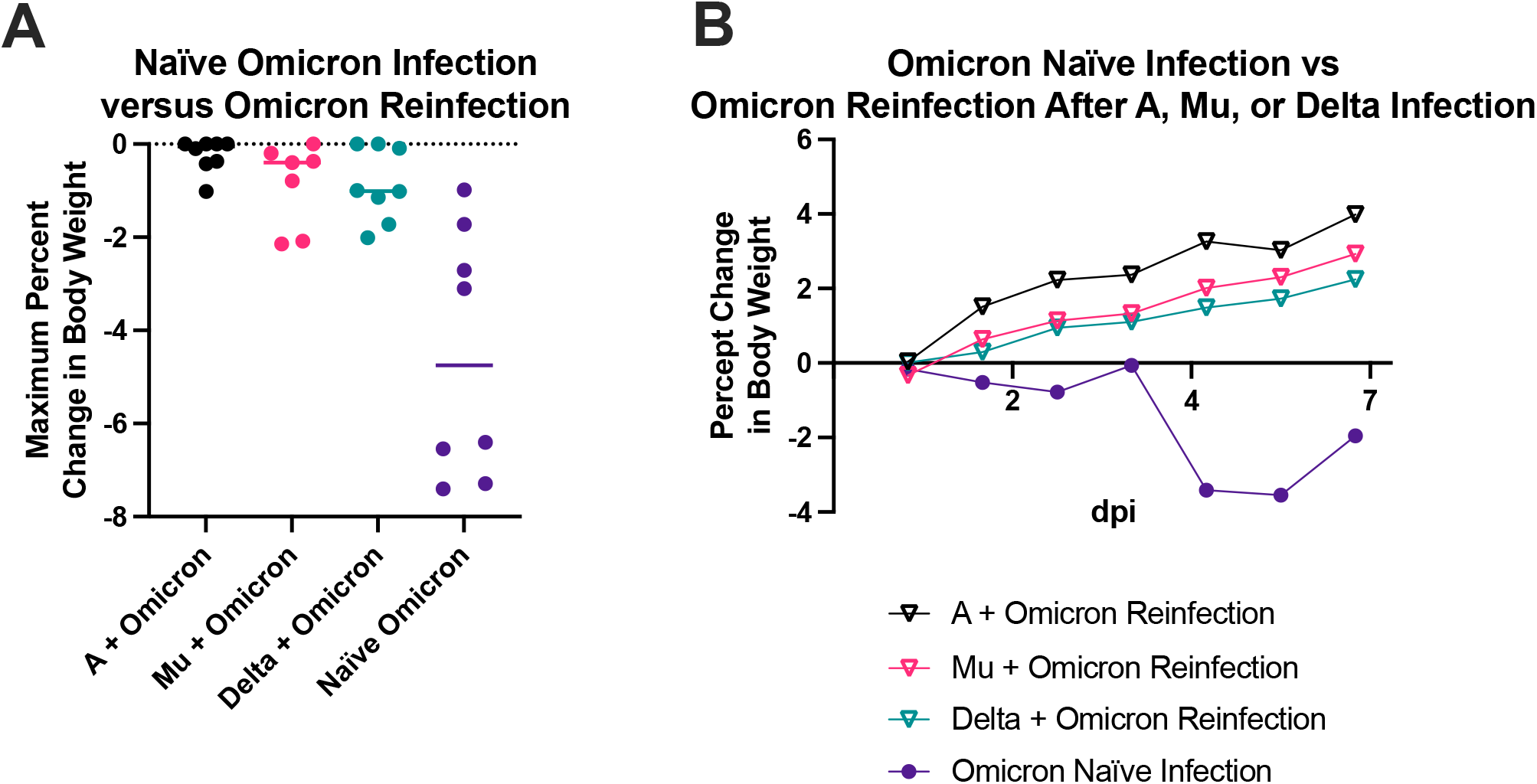
Percent change in body weight after naïve Omicron infection and Omicron reinfection. (A) Using a one-way ANOVA, the maximum percent change in body weight over 7 dpi was compared between naïve hamsters infected with Omicron, and hamsters reinfected with Omicron 28 days after initial infection with variant A, Mu, or Delta. There were no significant differences between reinfection groups, but all three reinfection groups had significantly less change in percent body weight compared to naïve Omicron infected hamsters (P<0.05). (B) The percent change in body weight is shown over 7 dpi with Omicron. Naïve hamsters infected with Omicron lost weight, while reinfected hamsters all gained weight on average regardless of their initial infection type.

**SUPPLEMENTAL TABLE 1.**
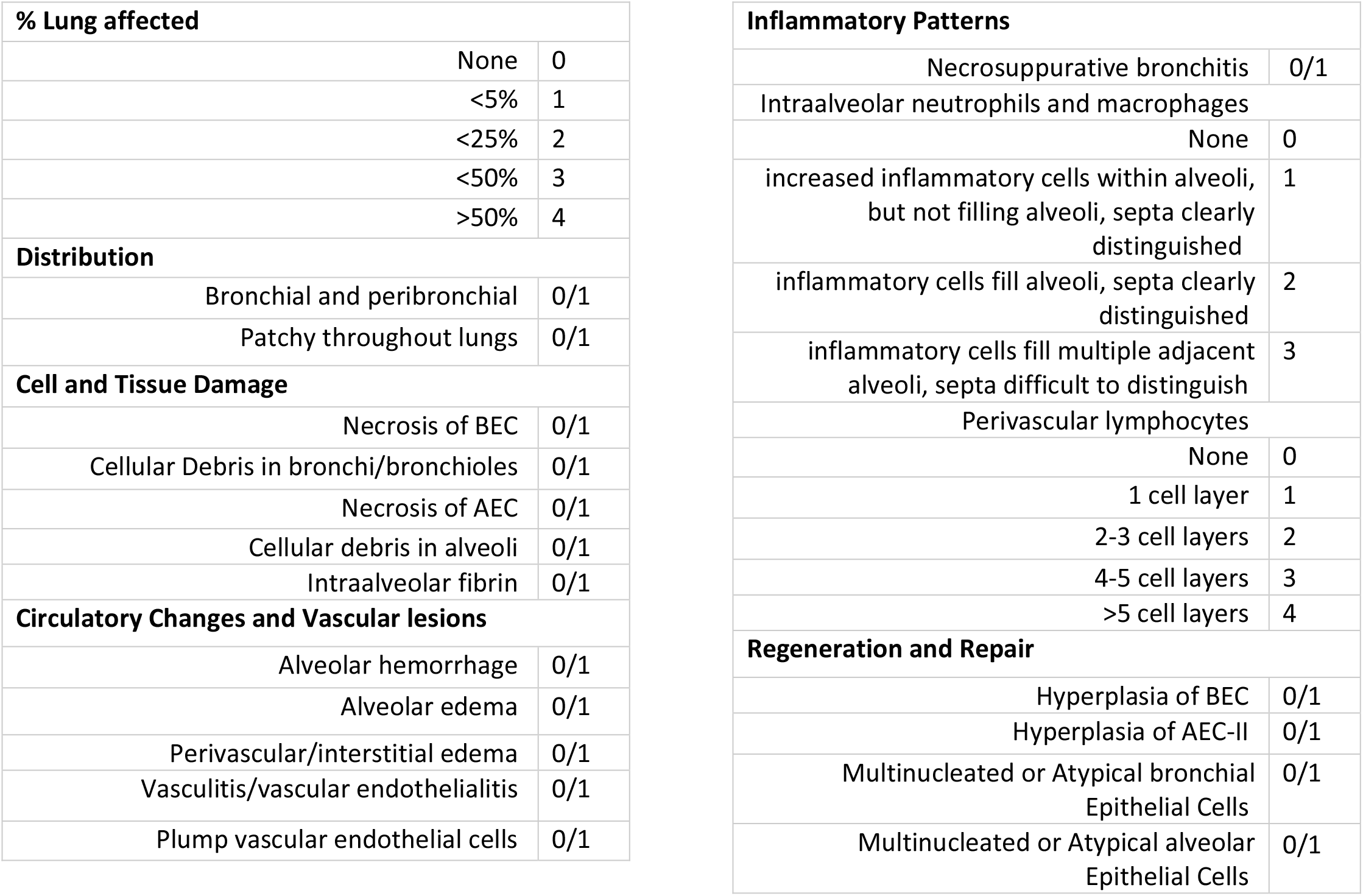
Histopathologic scoring system used by a blinded board-certified veterinary pathologist to analyze hamster lung tissue.

| % Lung affected |  |  | Inflammatory Patterns |  |
| --- | --- | --- | --- | --- |
|  | None | 0 | Necrosuppurative bronchitis | 0/1 |
|  | <5% | 1 | Intraalveolar neutrophils and macrophages |  |
|  | <25% | 2 | None | 0 |
|  | <50% | 3 | increased inflammatory cells within alveoli, but not filling alveoli, septa clearly distinguished | 1 |
|  | >50% | 4 | inflammatory cells fill alveoli, septa clearly distinguished | 2 |
| Distribution |  |  | inflammatory cells fill multiple adjacent alveoli, septa difficult to distinguish | 3 |
|  | Bronchial and peribronchial | 0/1 | Perivascular lymphocytes |  |
|  | Patchy throughout lungs | 0/1 | None | 0 |
| Cell and Tissue Damage |  |  | 1 cell layer | 1 |
|  | Necrosis of BEC | 0/1 | 2-3 cell layers | 2 |
|  | Cellular Debris in bronchi/bronchioles | 0/1 | 4-5 cell layers | 3 |
|  | Necrosis of AEC | 0/1 | >5 cell layers | 4 |
|  | Cellular debris in alveoli | 0/1 | Regeneration and Repair |  |
|  | Intraalveolar fibrin | 0/1 | Hyperplasia of BEC | 0/1 |
| Circulatory Changes and Vascular lesions |  |  | Hyperplasia of AEC-II | 0/1 |
|  | Alveolar hemorrhage | 0/1 | Multinucleated or Atypical bronchial Epithelial Cells | 0/1 |
|  | Alveolar edema | 0/1 | Multinucleated or Atypical alveolar Epithelial Cells | 0/1 |
|  | Perivascular/interstitial edema | 0/1 |  |  |
|  | Vasculitis/vascular endothelialitis | 0/1 |  |  |
|  | Plump vascular endothelial cells | 0/1 |  |  |

**SUPPLEMENTAL TABLE 2.**
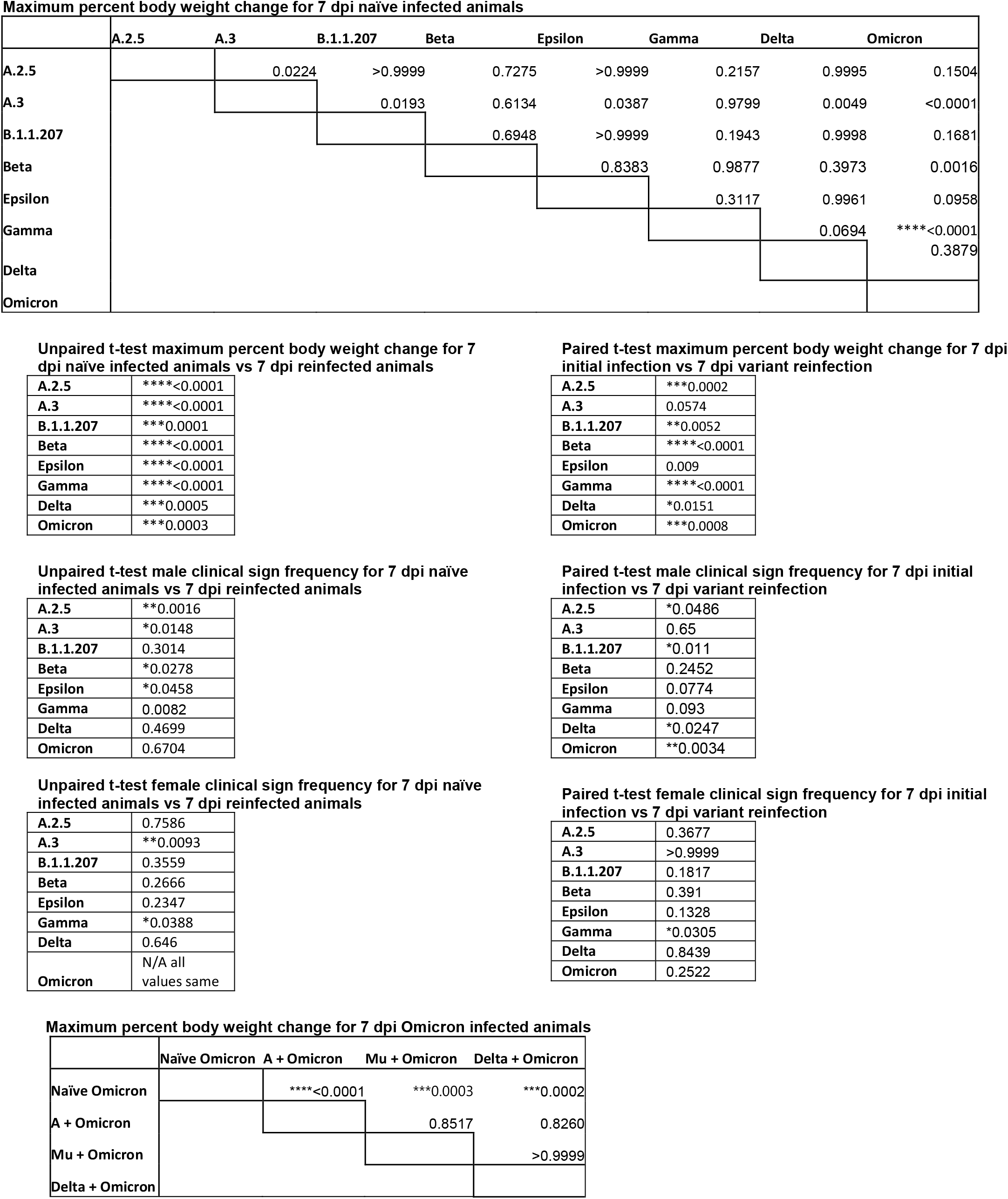
Results of multiple comparisons one-way ANOVA of maximum percent body weight change for 7 dpi naïve infected animals, unpaired T-test of maximum percent body weight change and clinical sign frequency for 7 dpi naïve infected animals vs 7 dpi reinfected animals, and paired T-test of maximum percent body weight change for 7 dpi initial infection vs 7 dpi variant reinfection.

**Maximum percent body weight change for 7 dpi naïve infected animals**
|  | A.2.5 | A.3 | B.1.1.207 | Beta | Epsilon | Gamma | Delta | Omicron |
| --- | --- | --- | --- | --- | --- | --- | --- | --- |
| A.2.5 |  | 0.0224 | >0.9999 | 0.7275 | >0.9999 | 0.2157 | 0.9995 | 0.1504 |
| A.3 |  |  | 0.0193 | 0.6134 | 0.0387 | 0.9799 | 0.0049 | <0.0001 |
| B.1.1.207 |  |  |  | 0.6948 | >0.9999 | 0.1943 | 0.9998 | 0.1681 |
| Beta |  |  |  |  | 0.8383 | 0.9877 | 0.3973 | 0.0016 |
| Epsilon |  |  |  |  |  | 0.3117 | 0.9961 | 0.0958 |
| Gamma |  |  |  |  |  |  | 0.0694 | ****<0.0001 |
| Delta |  |  |  |  |  |  |  | 0.3879 |
| Omicron |  |  |  |  |  |  |  |  |

**Unpaired t-test maximum percent body weight change for 7 dpi naïve infected animals vs 7 dpi reinfected animals**
|  |  |
| --- | --- |
| A.2.5 | ****<0.0001 |
| A.3 | ****<0.0001 |
| B.1.1.207 | ***0.0001 |
| Beta | ****<0.0001 |
| Epsilon | ****<0.0001 |
| Gamma | ****<0.0001 |
| Delta | ***0.0005 |
| Omicron | ***0.0003 |

**Paired t-test maximum percent body weight change for 7 dpi initial infection vs 7 dpi variant reinfection**
|  |  |
| --- | --- |
| A.2.5 | ***0.0002 |
| A.3 | 0.0574 |
| B.1.1.207 | **0.0052 |
| Beta | ****<0.0001 |
| Epsilon | 0.009 |
| Gamma | ****<0.0001 |
| Delta | *0.0151 |
| Omicron | ***0.0008 |

**Unpaired t-test male clinical sign frequency for 7 dpi naïve infected animals vs 7 dpi reinfected animals**
|  |  |
| --- | --- |
| A.2.5 | **0.0016 |
| A.3 | *0.0148 |
| B.1.1.207 | 0.3014 |
| Beta | *0.0278 |
| Epsilon | *0.0458 |
| Gamma | 0.0082 |
| Delta | 0.4699 |
| Omicron | 0.6704 |

**Paired t-test male clinical sign frequency for 7 dpi initial infection vs 7 dpi variant reinfection**
|  |  |
| --- | --- |
| A.2.5 | *0.0486 |
| A.3 | 0.65 |
| B.1.1.207 | *0.011 |
| Beta | 0.2452 |
| Epsilon | 0.0774 |
| Gamma | 0.093 |
| Delta | *0.0247 |
| Omicron | **0.0034 |

**Unpaired t-test female clinical sign frequency for 7 dpi naïve infected animals vs 7 dpi reinfected animals**
|  |  |
| --- | --- |
| A.2.5 | 0.7586 |
| A.3 | **0.0093 |
| B.1.1.207 | 0.3559 |
| Beta | 0.2666 |
| Epsilon | 0.2347 |
| Gamma | *0.0388 |
| Delta | 0.646 |
| Omicron | N/A all values same |

**Paired t-test female clinical sign frequency for 7 dpi initial infection vs 7 dpi variant reinfection**
|  |  |
| --- | --- |
| A.2.5 | 0.3677 |
| A.3 | >0.9999 |
| B.1.1.207 | 0.1817 |
| Beta | 0.391 |
| Epsilon | 0.1328 |
| Gamma | *0.0305 |
| Delta | 0.8439 |
| Omicron | 0.2522 |

|  | Naïve Omicron | A + Omicron | Mu + Omicron | Delta + Omicron |
| --- | --- | --- | --- | --- |
| Naïve Omicron |  | ****<0.0001 | ***0.0003 | ***0.0002 |
| A + Omicron |  |  | 0.8517 | 0.8260 |
| Mu + Omicron |  |  |  | >0.9999 |
| Delta + Omicron |  |  |  |  |

**SUPPLEMENTAL TABLE 3.**
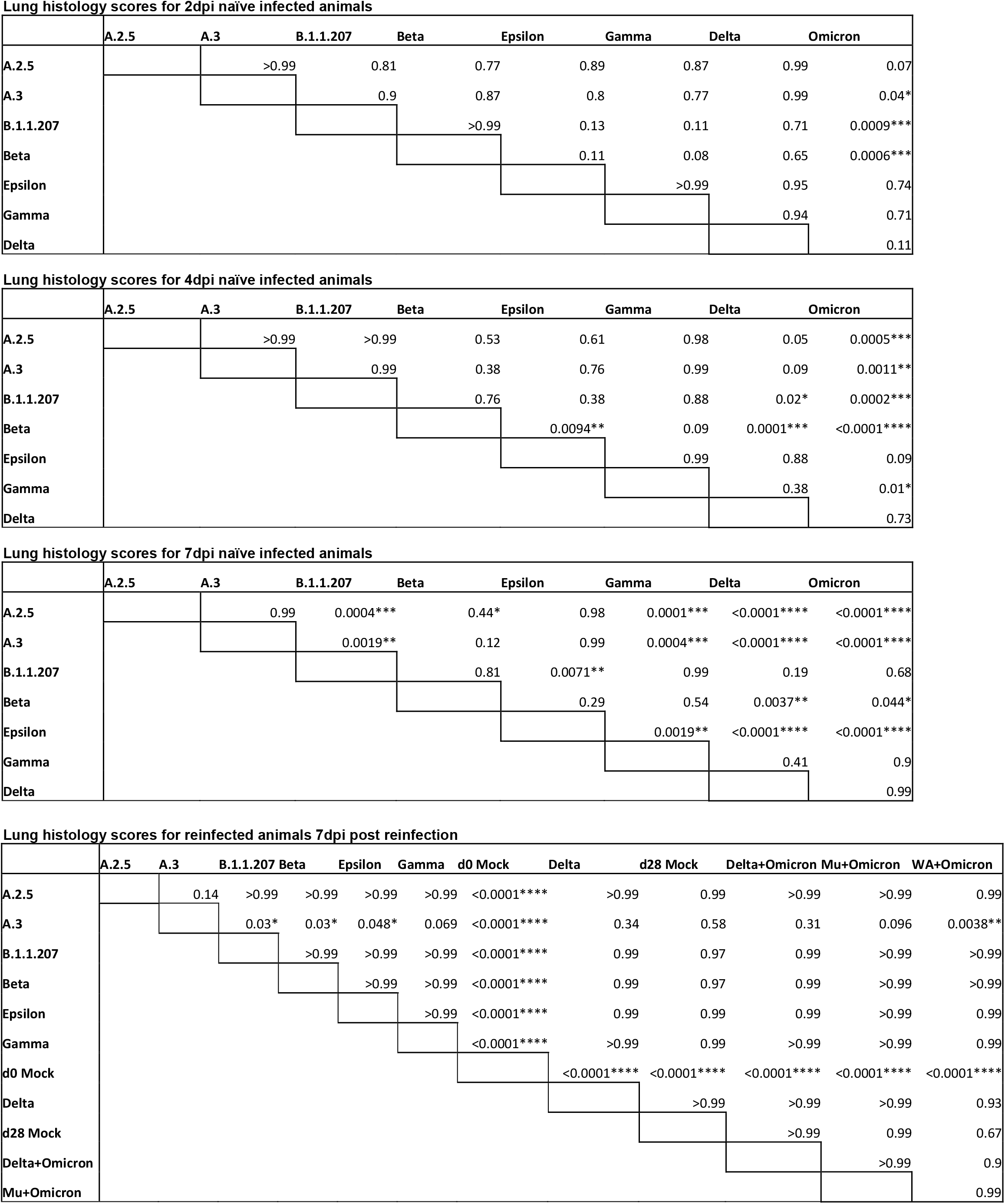
Results of multiple comparisons one-way ANOVA of histology scoring for 2, 4, and 7 dpi naïve infected animals and 7 dpi reinfected animals.

**SUPPLEMENTAL TABLE 4.** Results of multiple comparisons one-way ANOVA of viral titer for 2, 4, and 7 dpi naïve infected animals’ lung, trachea, and nasal turbinates.

|  | A.2.5 | A.3 | B.1.1.207 | Beta | Epsilon | Gamma | Delta | Omicron |
| --- | --- | --- | --- | --- | --- | --- | --- | --- |
| A.2.5 |  | ***0.0004 | >0.9999 | ****<0.0001 | 0.7891 | ****<0.0001 | 0.2977 | 0.9575 |
| A.3 |  |  | **0.0011 | 0.4592 | 0.0685 | 0.9991 | 0.3676 | ****<0.0001 |
| B.1.1.207 |  |  |  | ****<0.0001 | 0.9201 | ***0.0001 | 0.4801 | 0.8608 |
| Beta |  |  |  |  | ****<0.0001 | 0.8286 | **0.001 | ****<0.0001 |
| Epsilon |  |  |  |  |  | *0.0121 | 0.9943 | 0.1543 |
| Gamma |  |  |  |  |  |  | 0.1114 | ****<0.0001 |
| Delta |  |  |  |  |  |  |  | *0.0188 |
| Omicron |  |  |  |  |  |  |  |  |

**Lung viral titers 4dpi naïve infection**
|  | A.2.5 | A.3 | B.1.1.207 | Beta | Epsilon | Gamma | Delta | Omicron |
| --- | --- | --- | --- | --- | --- | --- | --- | --- |
| A.2.5 |  | 0.9958 | 0.8623 | *0.0232 | >0.9999 | ****<0.0001 | 0.6555 | ****<0.0001 |
| A.3 |  |  | 0.998 | **0.0019 | 0.9966 | ****<0.0001 | 0.21 | ****<0.0001 |
| B.1.1.207 |  |  |  | ***0.0001 | 0.8721 | ****<0.0001 | *0.0415 | ****<0.0001 |
| Beta |  |  |  |  | *0.0216 | 0.6263 | 0.7601 | ****<0.0001 |
| Epsilon |  |  |  |  |  | ****<0.0001 | 0.6405 | ****<0.0001 |
| Gamma |  |  |  |  |  |  | *0.0202 | ****<0.0001 |
| Delta |  |  |  |  |  |  |  | ****<0.0001 |
| Omicron |  |  |  |  |  |  |  |  |

**Lung viral titers 7dpi naïve infection**
|  | A.2.5 | A.3 | B.1.1.207 | Beta | Epsilon | Gamma | Delta | Omicron |
| --- | --- | --- | --- | --- | --- | --- | --- | --- |
| A.2.5 |  | >0.9999 | >0.9999 | >0.9999 | >0.9999 | >0.9999 | >0.9999 | >0.9999 |
| A.3 |  |  | >0.9999 | >0.9999 | >0.9999 | >0.9999 | >0.9999 | >0.9999 |
| B.1.1.207 |  |  |  | >0.9999 | >0.9999 | >0.9999 | >0.9999 | >0.9999 |
| Beta |  |  |  |  | >0.9999 | >0.9999 | >0.9999 | >0.9999 |
| Epsilon |  |  |  |  |  | >0.9999 | >0.9999 | >0.9999 |
| Gamma |  |  |  |  |  |  | >0.9999 | >0.9999 |
| Delta |  |  |  |  |  |  |  | >0.9999 |
| Omicron |  |  |  |  |  |  |  |  |

**Trachea viral titers 2dpi naïve infection**
|  | A.2.5 | A.3 | B.1.1.207 | Beta | Epsilon | Gamma | Delta | Omicron |
| --- | --- | --- | --- | --- | --- | --- | --- | --- |
| A.2.5 |  | 0.0535 | 0.5398 | 0.7059 | >0.9999 | 0.8984 | ****<0.0001 | **0.0037 |
| A.3 |  |  | ****<0.0001 | 0.8678 | 0.1515 | 0.6567 | *0.0112 | ****<0.0001 |
| B.1.1.207 |  |  |  | *0.0101 | 0.2833 | *0.0326 | ****<0.0001 | 0.5128 |
| Beta |  |  |  |  | 0.9123 | >0.9999 | ****<0.0001 | ****<0.0001 |
| Epsilon |  |  |  |  |  | 0.9869 | ****<0.0001 | ***0.0008 |
| Gamma |  |  |  |  |  |  | ****<0.0001 | ****<0.0001 |
| Delta |  |  |  |  |  |  |  | ****<0.0001 |
| Omicron |  |  |  |  |  |  |  |  |

Trachea viral titers 4dpi naïve infection
|  | A.2.5 | A.3 | B.1.1.207 | Beta | Epsilon | Gamma | Delta | Omicron |
| --- | --- | --- | --- | --- | --- | --- | --- | --- |
| A.2.5 |  | >0.9999 | 0.1954 | 0.9987 | 0.4417 | >0.9999 | 0.8416 | 0.5052 |
| A.3 |  |  | 0.2246 | 0.9994 | 0.4863 | >0.9999 | 0.8722 | 0.551 |
| B.1.1.207 |  |  |  | 0.5499 | 0.9998 | 0.1799 | 0.9589 | 0.9994 |
| Beta |  |  |  |  | 0.8312 | 0.998 | 0.9919 | 0.8752 |
| Epsilon |  |  |  |  |  | 0.4167 | 0.9983 | >0.9999 |
| Gamma |  |  |  |  |  |  | 0.8224 | 0.4793 |
| Delta |  |  |  |  |  |  |  | 0.9994 |
| Omicron |  |  |  |  |  |  |  |  |

Nasal Turbinates viral titers 2dpi naïve infection
|  | A.2.5 | A.3 | B.1.1.207 | Beta | Epsilon | Gamma | Delta | Omicron |
| --- | --- | --- | --- | --- | --- | --- | --- | --- |
| A.2.5 |  | 0.2694 | >0.9999 | 0.2075 | 0.9616 | 0.2079 | 0.9954 | 0.2482 |
| A.3 |  |  | 0.1613 | >0.9999 | 0.9033 | >0.9999 | 0.7431 | ***0.0001 |
| B.1.1.207 |  |  |  | 0.119 | 0.8899 | 0.1193 | 0.9747 | 0.3862 |
| Beta |  |  |  |  | 0.8487 | >0.9999 | 0.6597 | ****<0.0001 |
| Epsilon |  |  |  |  |  | 0.8493 | >0.9999 | *0.0148 |
| Gamma |  |  |  |  |  |  | 0.6604 | ****<0.0001 |
| Delta |  |  |  |  |  |  |  | *0.0401 |
| Omicron |  |  |  |  |  |  |  |  |

Nasal Turbinates viral titers 4dpi naïve infection
|  | A.2.5 | A.3 | B.1.1.207 | Beta | Epsilon | Gamma | Delta | Omicron |
| --- | --- | --- | --- | --- | --- | --- | --- | --- |
| A.2.5 |  | 0.5567 | 0.7599 | 0.9997 | 0.7113 | 0.3313 | 0.9231 | ***0.0002 |
| A.3 |  |  | *0.0146 | 0.2606 | *0.0112 | >0.9999 | 0.9977 | ****<0.0001 |
| B.1.1.207 |  |  |  | 0.9555 | >0.9999 | **0.0044 | 0.0997 | 0.0517 |
| Beta |  |  |  |  | 0.9361 | 0.1225 | 0.683 | **0.0013 |
| Epsilon |  |  |  |  |  | **0.0033 | 0.0814 | 0.0643 |
| Gamma |  |  |  |  |  |  | 0.9712 | ****<0.0001 |
| Delta |  |  |  |  |  |  |  | ****<0.0001 |
| Omicron |  |  |  |  |  |  |  |  |

Nasal Turbinates viral titers 7dpi naïve infection
|  | A.2.5 | A.3 | B.1.1.207 | Beta | Epsilon | Gamma | Delta | Omicron |
| --- | --- | --- | --- | --- | --- | --- | --- | --- |
| A.2.5 |  | 0.9485 | 0.9554 | 0.9916 | >0.9999 | 0.2087 | 0.9998 | >0.9999 |
| A.3 |  |  | >0.9999 | >0.9999 | 0.9881 | 0.877 | 0.9972 | 0.9485 |
| B.1.1.207 |  |  |  | >0.9999 | 0.9904 | 0.8639 | 0.9979 | 0.9554 |
| Beta |  |  |  |  | 0.9992 | 0.7112 | >0.9999 | 0.9916 |
| Epsilon |  |  |  |  |  | 0.3425 | >0.9999 | >0.9999 |
| Gamma |  |  |  |  |  |  | 0.4569 | 0.2087 |
| Delta |  |  |  |  |  |  |  | 0.9998 |
| Omicron |  |  |  |  |  |  |  |  |

